## Supplemental Data for "MKP1 promotes nonalcoholic steatohepatitis by suppressing AMPK activity through LKB1 nuclear retention"

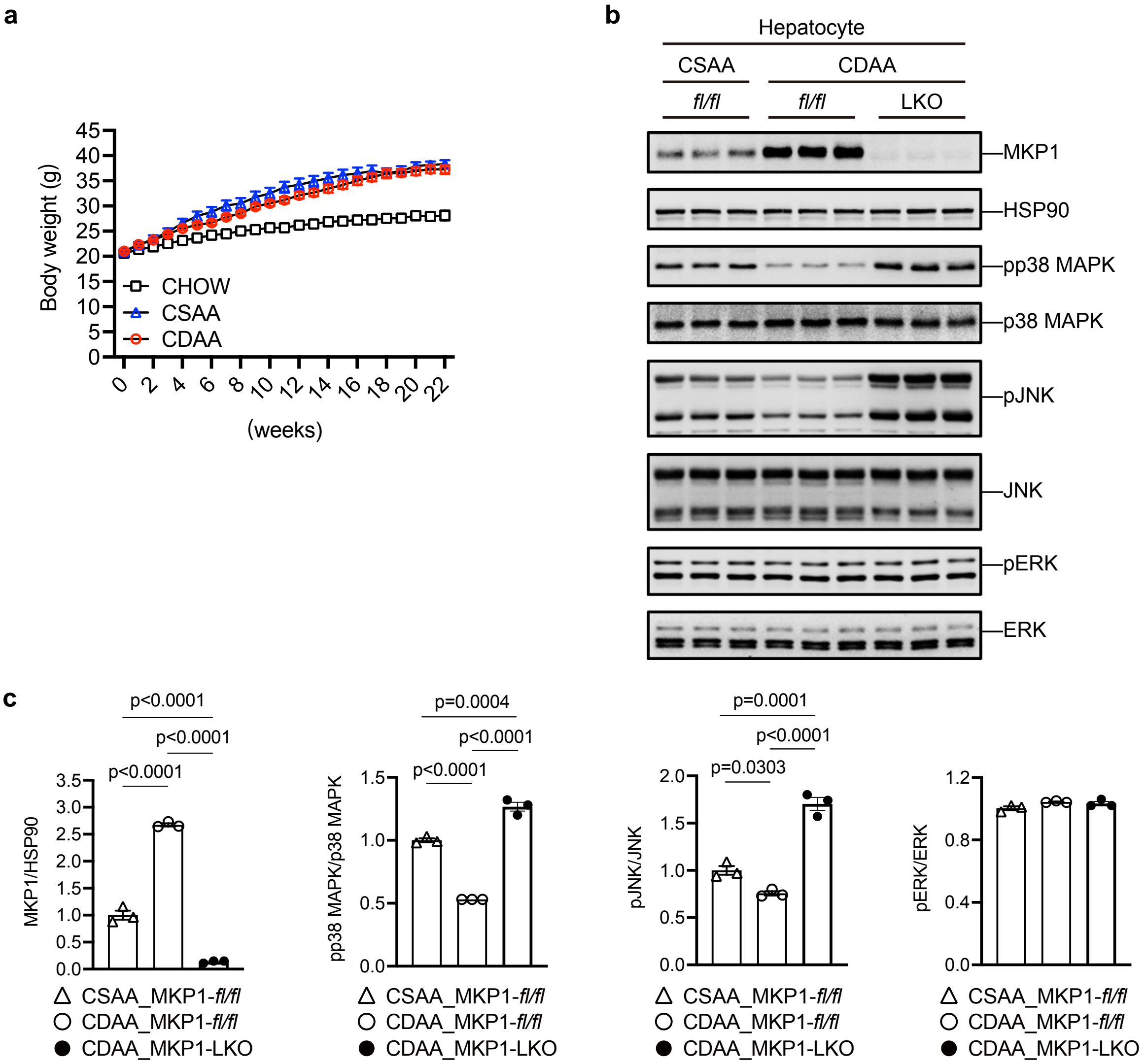


**Supplemental Figure 1. MKP1 expression and MAPK regulation in hepatocytes from NASH diet fed mice.** (**a**) Growth curve from male *Mkp1*$^{fl/fl}$ mice fed with Chow, CSAA or CDAA diet for 22 weeks. The body weight was recorded weekly. Data represent the mean ± SEM from 10-13 mice for each diet. (**b**) Immunoblots of MKP1 and HSP90 as a loading control in hepatocytes and p38 MAPK, pJNK and pERK with corresponding MAPK totals from isolated hepatocytes derived from male *Mkp1*$^{fl/fl}$ and MKP1-LKO mice fed with CSAA or CDAA diet for 8 weeks. (**c**) Quantitation of immunoblots from (**b**). Key: *fl/fl, Mkp1*$^{fl/fl}$; LKO, MKP1-LKO. Data represent the mean ± SEM from 3 mice per group. *p* values were determined by one-way ANOVA.

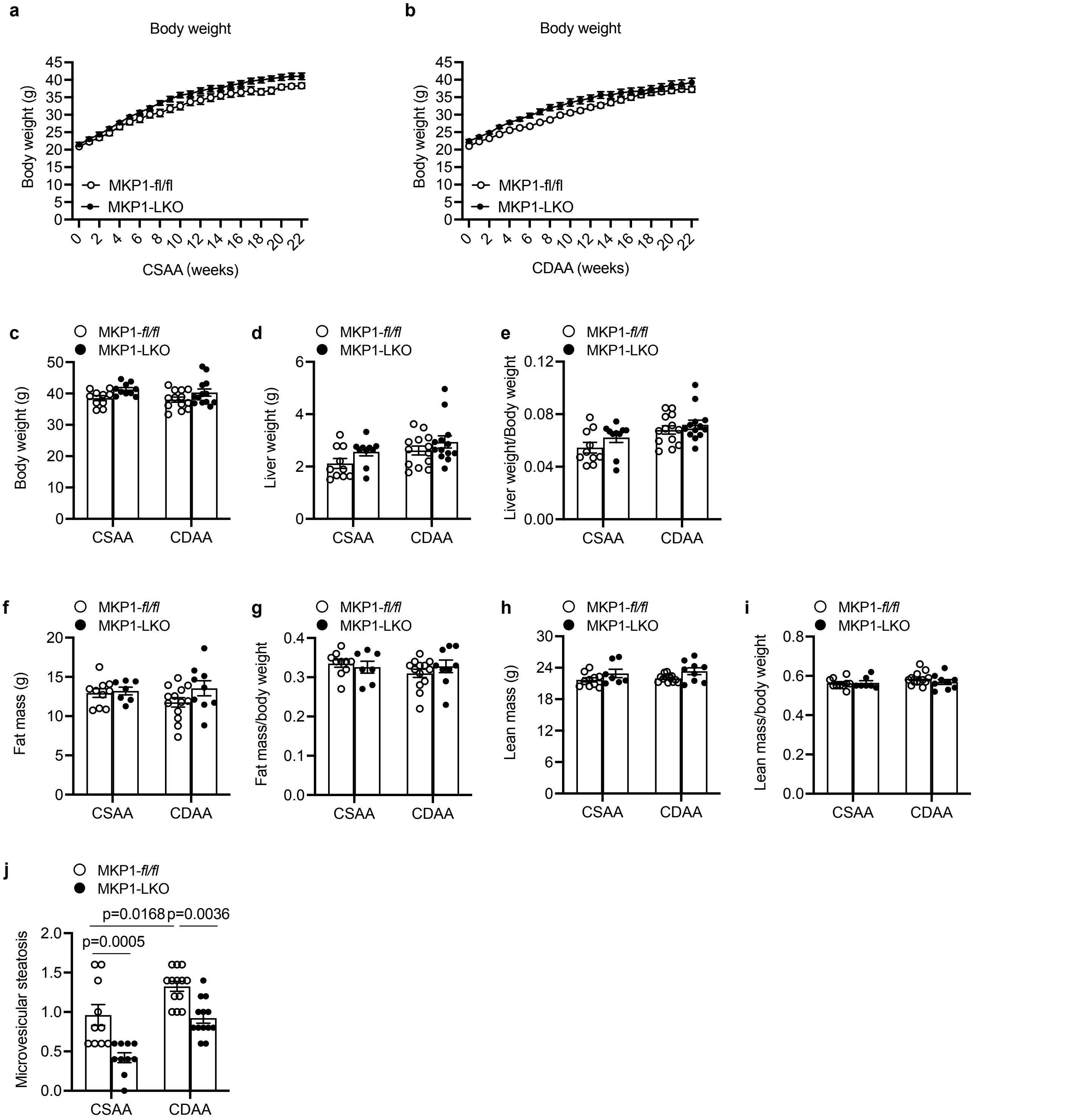


**Supplemental Figure 2. Characterization of *Mkp1$^{fl/fl}$* and MKP1-LKO mice fed with CSAA or CDAA diet.** (**a** and **b**) Growth curve from male *Mkp1$^{fl/fl}$* and MKP1-LKO mice fed with CSAA or CDAA diet for 22 weeks, respectively and body weight was recorded weekly. (**c-e**) Body weight, liver weight and ratio of liver/body weight after 22 weeks of CSSAA and CDAA diet feeding. Data represent the mean ± SEM from 10-13 mice for each diet. *p* values shown were determined by two-way ANOVA. (**f-i**) The body composition in *Mkp1$^{fl/fl}$* and MKP1-LKO mice fed with CSAA or CDAA diet for 22 weeks, presented as (**f**) fat mass, (**g**) fat mass/body weight, (**h**) lean mass and (**i**) lean mass/body weight. (**j**) Quantification of microvesicular steatosis in **Fig. 1c**. Data represent mean ± SEM from 7-13 mice for each diet. *p* values shown in (**c**), (**f-j**) were determined by two-way ANOVA, shown in (**d**) and (**e**) were determined by Kruskal-Wallis test.

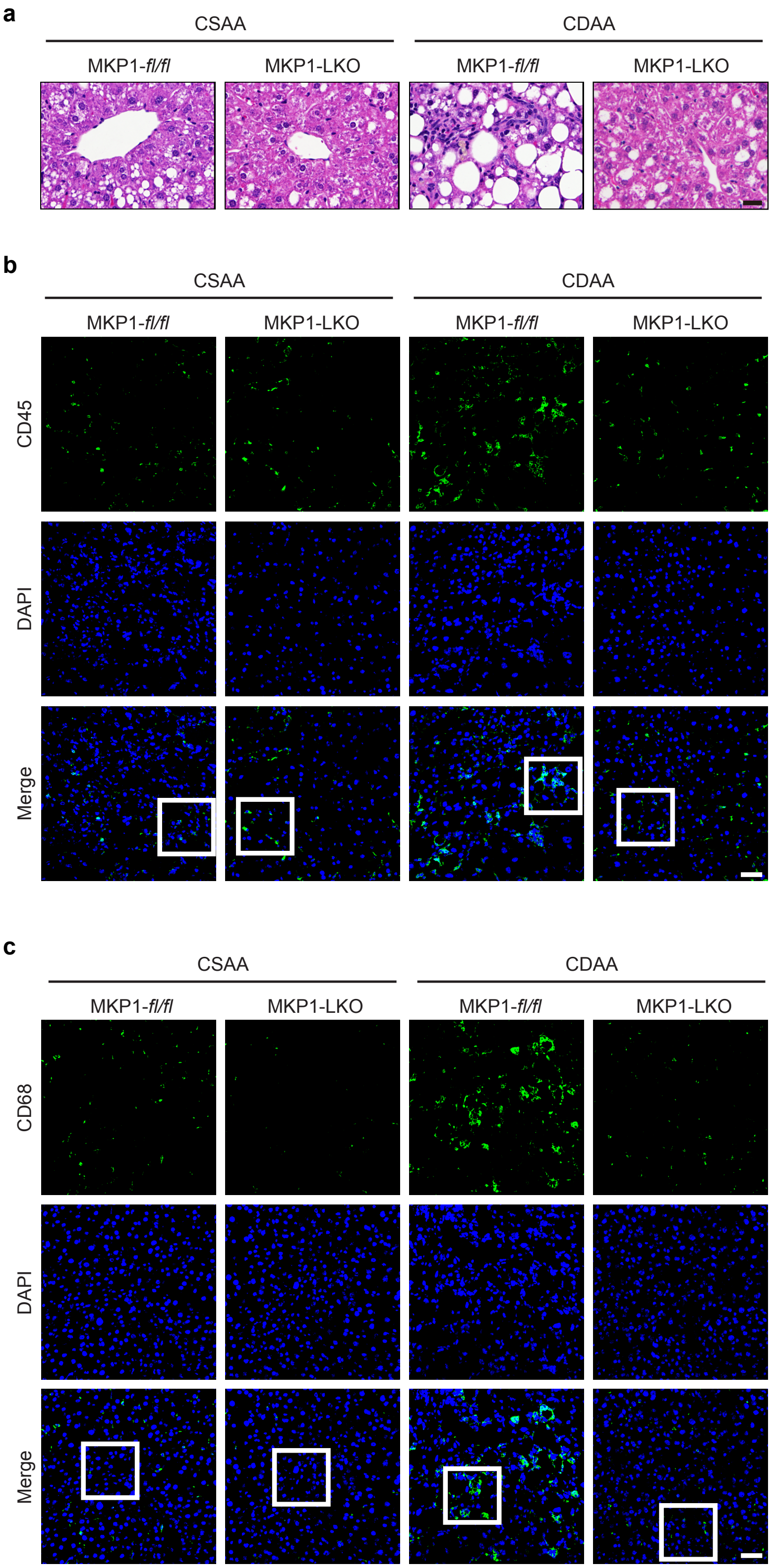


**Supplemental Figure 3. Histological analyses of inflammatory infiltrates in MKP1-LKO mice fed with a NASH diet.** *Mkp1*$^{fl/fl}$ and MKP1-LKO mice were fed with CSAA or CDAA diet for 22 weeks. (**a**) Histological examination of inflammatory infiltrates in livers. Scale bar = 25 μm. (**b** and **c**) Staining of CD45 or CD68 in liver sections from CSAA or CDAA fed *Mkp1*$^{fl/fl}$ and MKP1-LKO mice. Scale bar = 50 μm. High power magnification of the indicated rectangle areas are shown in **Fig. 3a** and **3d**.

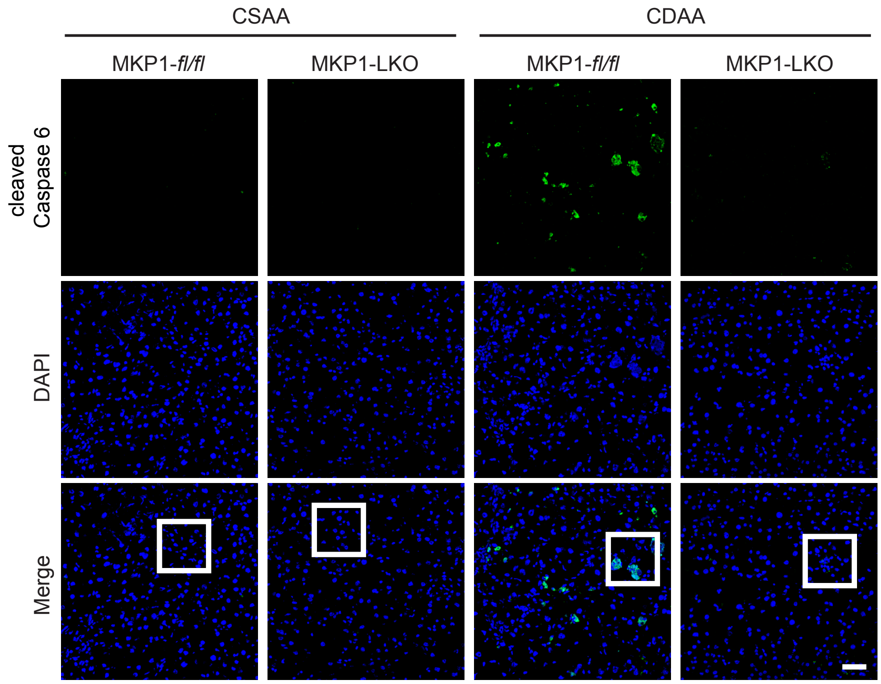


**Supplemental Figure 4. MKP1 promotes caspase 6 cleavage in NASH-diet fed mice.** Staining of cleaved caspase 6 in liver sections from *Mkp1*$^{fl/fl}$ and MKP1-LKO mice fed with CSAA or CDAA diet for 22 weeks. Scale bar = 50 µm. High power magnification of the indicated rectangle areas are shown in **Fig. 4l**.

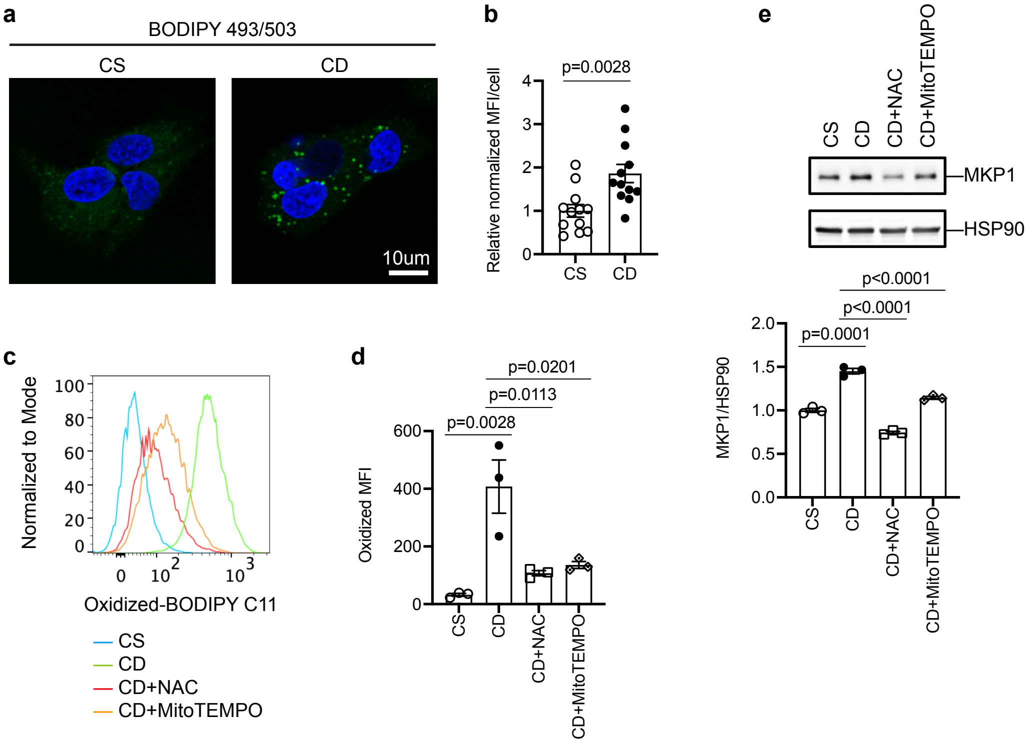


**Supplemental Figure 5. MKP1 is upregulated in a ROS-dependent manner in liver cells.** HepG2 cells were treated with choline-sufficient (CS) or choline-deficient (CD) medium in the absence or presence of 2 mM NAC or 20 µM MitoTEMPO after 24 h starvation in serum-free medium. (**a**) Lipid content was detected using BODIPY 493/503 (Green) and cell nuclei stained with DAPI (Blue). Scale bar = 10 µm. **(b)** The mean fluorescence intensity (MFI) per field was quantified using ImageJ with each field containing between 2-9 cells. A total of 12 fields were analyzed for each condition (48 cells for CS treated cells and 51 cells for CD treated cells), and MFI values were normalized on a per-cell basis. (**c**) ROS content was measured using BODIPY 581/591 C11. (**d**) MFI signals were calculated by FlowJo. Data represent the mean ± SEM from 3 independent experiments. *p* values were determined by student's-unpaired t test. (**e**) Immunoblots of MKP1 and HSP90 as a loading control. Lower panel represents the quantitation of immunoblots. Data represent the mean ± SEM from 3 independent experiments. *p* values shown in (**b**) were determined by student's-unpaired t test, shown in (**d**) and (**d**) were determined by one-way ANOVA. Key: CS, choline sufficient medium; CD, choline-deficient medium; NAC, N-acetyl-l-cysteine; MFI, mean fluorescence intensity.

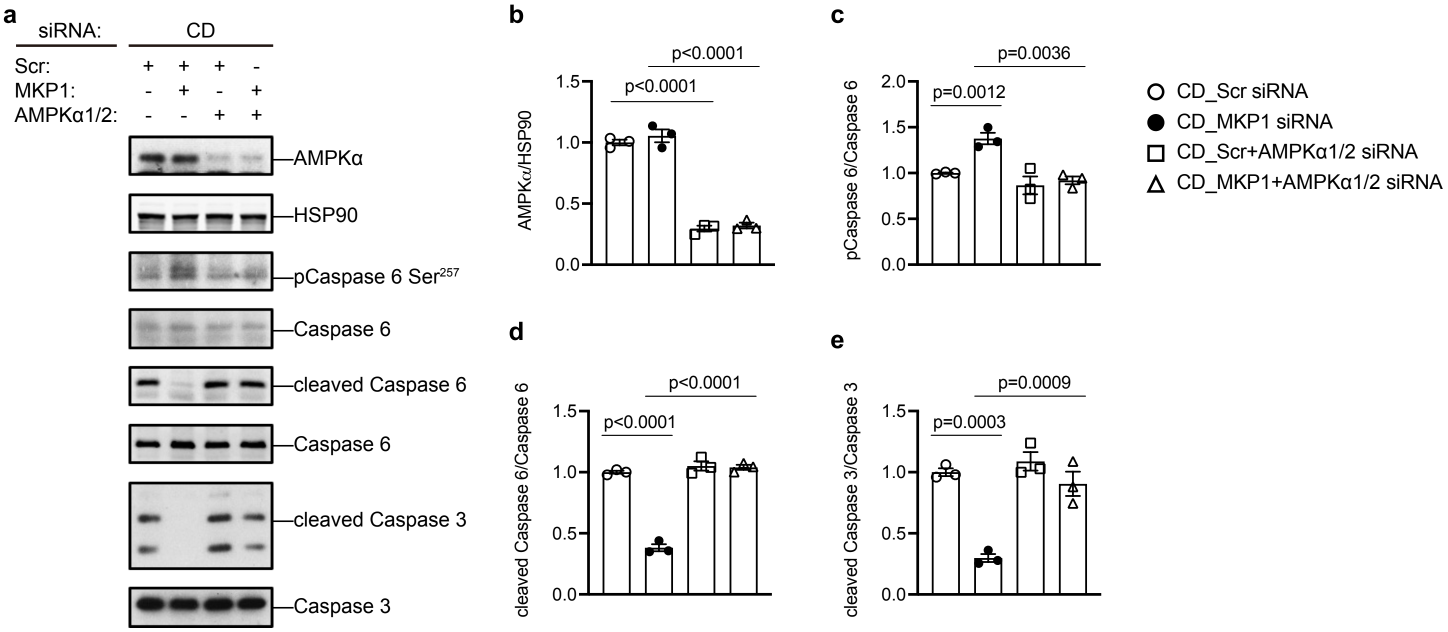


**Supplemental Figure 6. MKP1 acts upstream of the AMPKα-caspase 6 pathway.** Knockdown of AMPKα1/2 rescues the effects of MKP1 deficiency on Caspase 6/3 cleavage. (**a**) Immunoblots of AMPKα, phospho-caspase 6 (Ser257), cleaved Caspase 6 and cleaved caspase 3 with the indicated corresponding totals. (**b-e**) Quantitation of immunoblots from (**a**). Key: Scr, scrambled siRNA; MKP1, MKP1 siRNA; AMPKα1/2, AMPKα1 siRNA+ AMPKα2 siRNA; CD, choline-deficient medium. Data represent the mean ± SEM from 3 independent experiments. *p* values were determined by two-way ANOVA.

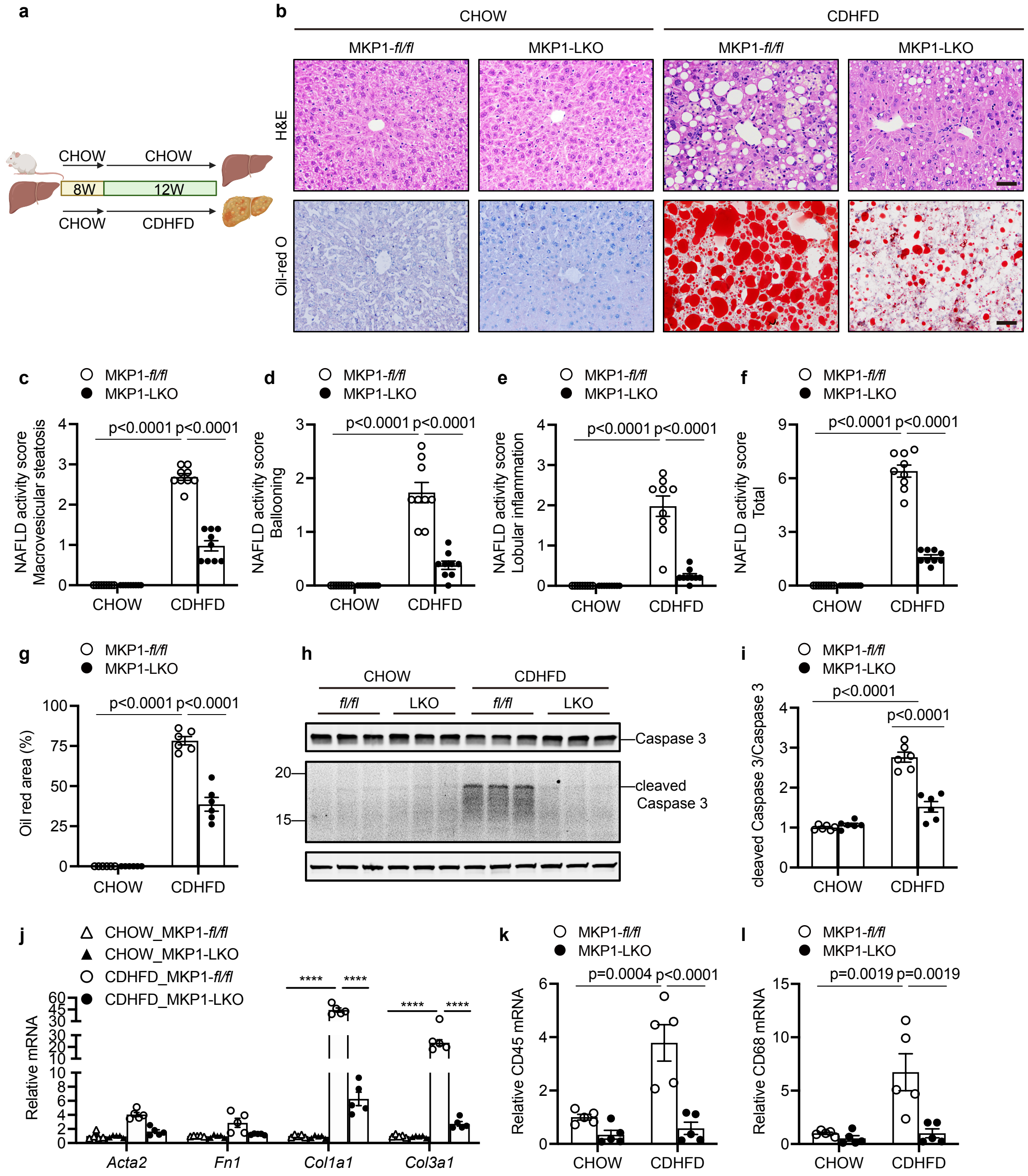


**Supplemental Figure 7. Loss of hepatic MKP1 prevents CDHFD diet-induced NASH.** The *Mkp1*$^{fl/fl}$ and MKP1-LKO mice were fed with either Chow or CDHFD (60% kcal fat) diet for 12 weeks. (**a**) Schematic diagram for Chow/CDHFD diet. Key: 8W, 8 weeks; 12W, 12 weeks. (**b**) Histological H&E staining (Scale bar = 50 μm) and Oil red-O staining (Scale bar = 50 μm) from liver sections. (**c-f**) NAFLD activity score for macrovesicular steatosis (**c**), ballooning (**d**), lobular inflammation (**e**) and total NAFLD score (**f**) from H&E staining in (**b**). Data represent the mean ± SEM derived from 9-10 mice per genotype. (**g**) Quantification of Oil red O-stained areas from Oil red-O staining in (**b**). Data represent the mean ± SEM derived from 6 mice per group. (**h**) Immunoblot of expression of cleaved caspase 3 and caspase 3 in livers. (**i**) Densitometry of immunoblots from cleaved caspase 3/caspase 3 from (**h**). Data represent the mean ± SEM from 6 mice per group. Key: *fl/fl, Mkp1*$^{fl/fl}$; LKO, MKP1-LKO. (**j**) mRNA expression of fibrotic genes in livers of mice fed chow or CDHFD for 8 weeks. Data represent the mean ± SEM derived from 5 mice per group. (**k** and **l**) mRNA expression of CD45 and CD68 in livers from Chow or CDHFD fed *Mkp1*$^{fl/fl}$ and MKP1-LKO mice. Data represent the mean ± SEM derived from 5 mice per group. *p* values shown in (**i-l**) were determined by two-way ANOVA, *p* values shown in (**c-f**) were determined by Kruskal-Wallis test.

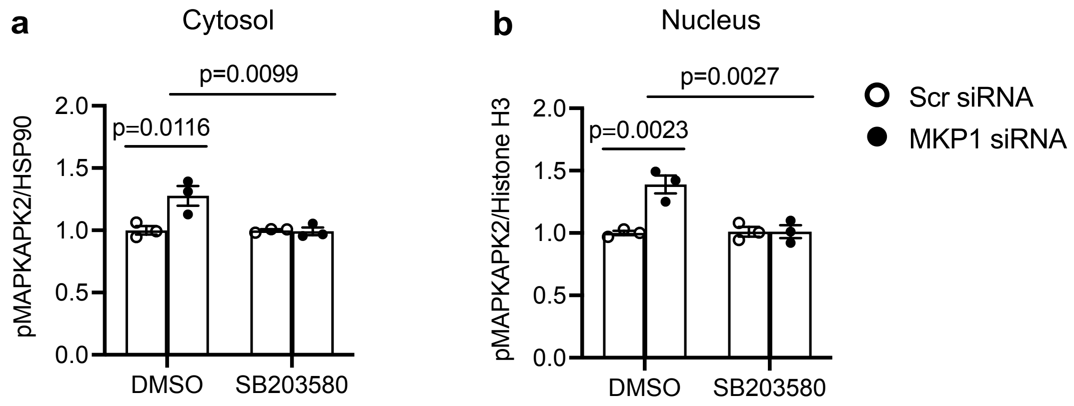


**Supplemental Figure 8. MKP1-mediated p38 MAPK activity in SB203580-treated HepG2 cells.** Densitometry as a ratio of phospho-MAPKAPK2 (Thr334)/ HSP90 in the cytosol (**a**) and nucleus (**b**) were from immunoblots in **Figure 9a**. Data represent mean ± SEM from 3 independent experiments. *p* values were determined by two-way ANOVA.

### Supplemental Figure 9

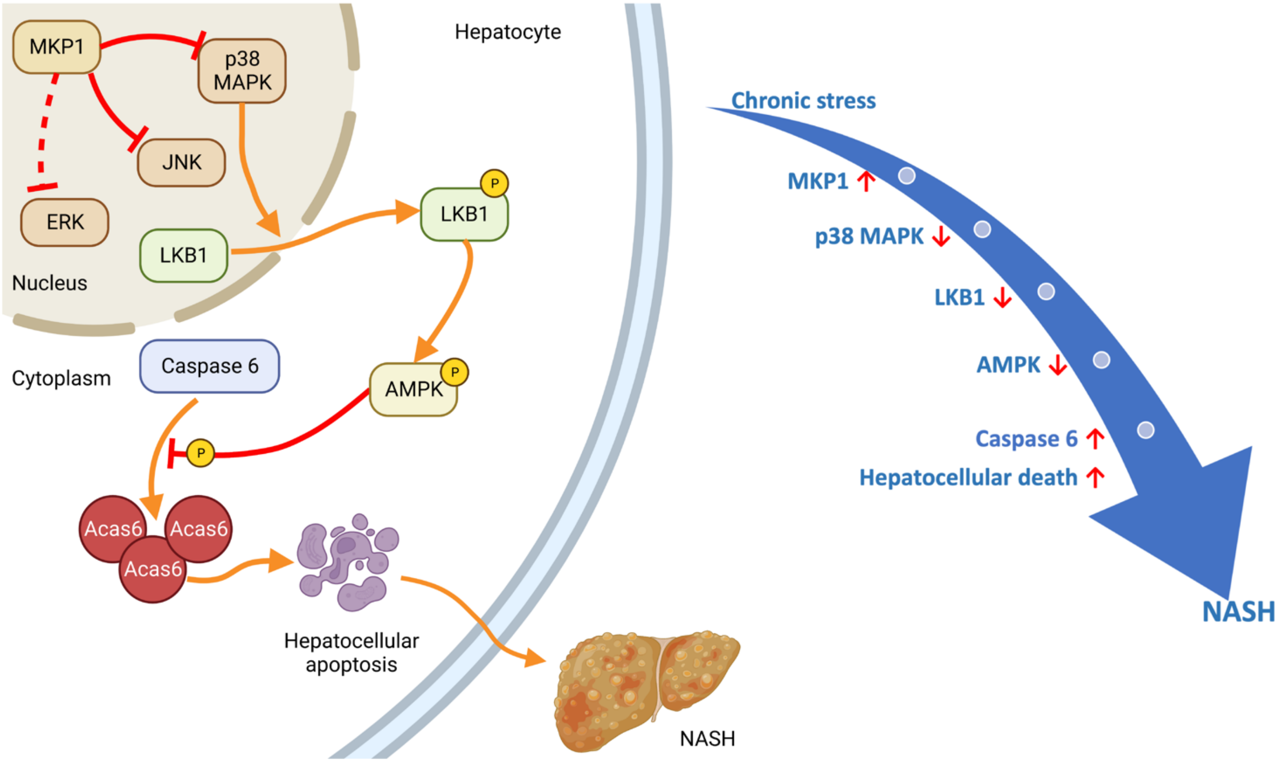


**Supplemental Figure 9. Model for the regulation MKP1-p38 MAPK-AMPK-Caspase 6 pathway in NASH.** See text for details.
